# Identification of whole-body reaching movement phenotypes in young and older active adults: an unsupervised machine learning approach

**DOI:** 10.1101/2024.07.09.600023

**Authors:** Michel Pfaff, Matthieu Casteran

## Abstract

Studies reported age-related motor control modifications in whole-body movement in several aspects of spatiotemporal movement organization by comparing young and older adults. However, studies on motor control involve high complexity and high-dimensional data of different natures, in which machine learning has proved to be effective. Furthermore, conventional studies focus on comparisons of movement parameters based on a priori grouping, whereas unsupervised machine learning allows the identification of inherent groupings within the dataset. The current investigation was carried out by using the unsupervised machine learning on motor control features across age-groups. An important question was whether we could identify different movement patterns based on motor control features and whether they were age-dependent or independent. We investigated motor control parameters variations in a whole-body reaching movement across young and active older adults including woman and man (n=19). We applied the K-means clustering algorithm to segment the kinematic data (21 features) of all individuals. We propose a methodology applying the latest recommendations for clustering methods in the field of whole-body movement motor control. Analysis revealed two distinct motor control patterns which were age independent. The first pattern exhibited higher shoulder, ankle and knee angular excursions, along with a higher vertical velocity of center of mass (CoM), compared to the second pattern, which had higher hip and back angular excursions, along with a lower vertical velocity CoM. The clustering methodology demonstrated its effectiveness to identify distinct motor patterns based solely on motor control features independently of age-grouping.

**Significance Statement:**

- K-means clustering algorithm enabled us to identify two distinct age-independent motor patterns: a first pattern with high shoulder, ankle and knee angular excursions, and vertical velocity of CoM; a second pattern with high hip and back angular excursions and low vertical velocity of CoM.
- Demonstrates how unsupervised machine learning can identify motor patterns and proposes a methodology to apply it in the field of whole-body movement motor control.
- Proves the complementary contribution of unsupervised machine learning to conventional approach for motor control studies, which enables to process the high complexity and dimensionality of movements.
- Advances understanding of motor behaviours through unsupervised machine learning analysis of whole-body reaching movements.

## 1. Introduction

Human motor control was investigated under various dimensions, such as the number of segment involved, velocity, based of support and constraint of movement, and across age (Alexandrov et al., 2001; Thomas et al., 2005; Berret et al., 2009; Fautrelle et al., 2010; Casteran et al., 2013, 2018). Specifically, the latter focuses on differences between young and older adults and reveals significant insights into the aging process of motor control. (Konrad et al., 1999; Paizis et al., 2008; Bilodeau-Mercure et al., 2015; Cruz-Jimenez, 2017; Larivière et al., 2019). Specifically, whole-body movement studies show age-related differences in several aspects of spatiotemporal movement organization (Patla et al., 1993; Mourey et al., 2000; Paizis et al., 2008; Casteran et al., 2018; Honda et al., 2021, 2022) and age-related motor patterns (Papa and Cappozzo, 2000; Casteran et al., 2018).

Studies on motor control involve high complexity and high-dimensional data of different natures. Due to motor control complexity, several authors are interested in advanced technologies, such as unsupervised and supervised machine learning, to study human movements (Halilaj et al., 2018), particularly in the study of motor control across age (Ackermans et al., 2019; Guo et al., 2022; Liang et al., 2022). The contribution of machine learning in motor control studies becomes relevant to improve and complement conventional analysis methods. In fact, the kinematic data characterising human movement is highly dimensional and often heterogeneous as shown by Schwarz et al. (2019) which identified over 50 kinematic parameters for reaching movements in elderly stroke patient. In addition, current data from motor control studies differ in nature (kinematic, kinetic, and physiological) and are high resolution requiring large-scale, different and parallel processing. The contribution of unsupervised machine learning enables processing, based on groups inherent in the dataset, without prior matching of groups.

Recent literature underscores that unsupervised machine learning models such as clustering techniques like K-means, are able to identify different patterns of movements based on kinematic data (Arac, 2020). For instance, studies by Ackermans et al. (2019) and Sawacha et al. (2020) have used clustering methods to segment kinematic and kinetic data, demonstrating its complementary relevance in motor control research. Moreover, recent work by Higgins et al. (2023) applies these methods to analyse kinematic time-series data of lumbar and pelvic movements, while Liang et al. (2022) use them to distinguish consistent gait characteristics, thereby aiding in the assessment of normal gait functions. Guo et al. (2022) explore age-related changes in muscle synergies and their activations during walking. Nevertheless, the effective deployment of machine learning within studies on motor control necessitates adherence to established best practices to ensure reliability and efficacy. The review of Halilaj et al. (2018) summarizes the current usage of machine learning methods in human movement biomechanics and highlights best practices that will enable critical evaluation of the literature. Furthermore, the study by Zakharov (2016) provides a comprehensive overview of K-means clustering analysis, aiming to enhance the methodological clarity and the quality of future research.

The current investigation was carried out by using the K-means algorithm to identify movement profiles in the kinematic features of whole body reaching (WBR) movements including different age groups (young and older). The purpose of the study was to determine whether K-means could identify different movement patterns based on motor control features and whether they were age-dependent or independent. We hypothesize that the clustering methodology will identify at least two clusters (age or motor pattern related) based on the current motor control literature. We propose a methodology for the K-means algorithm including current best practices, to detect motor patterns based on kinematic features during a WBR movements.

## 2. Material and methods

### 2.1. Subject

Data for this study were extracted from the study of Casteran et al. (2013 and 2018), specifically the natural speed and 30% target conditions. The young subjects were 10 healthy adults: six women and four men (mean age: 24 ± 2 years; mean height: 170 ± 0.08 m; mean weight: 59 ± 11 kg). In the case of the healthy aging subjects, there were five women and four men (mean age: 70 ± 2 years; mean height: 163 ± 0.09 m; mean weight: 62 ± 13 kg). Subjects were excluded in cases of falls, prostheses, neurological diseases, a history of alcohol abuse, taking drugs affecting central nervous system functions, genetic metabolic diseases, diabetes, hypertension and hypotension. They had normal or corrected to normal vision. The physical fitness of the aging subjects was evaluated using the Timed Up and Go test (TUG (Podsiadlo and Richardson, 1991)), the Unipodal Balance Test (UBT (Hurvitz et al., 2000)), and the Five-Times-Sit-to-Stand Test (FTSST adapted to Csuka and McCarty (Csuka and McCarty, 1985; Guralnik et al., 1994)). The subjects performed the TUG test below 8 seconds indicating no mobility problems (Lusardi et al., 2003). They were able to stay more than 30 seconds in a unipodal position hence indicating a very low risk of falling (Hurvitz et al., 2000). They were all able to achieve their FTSST in less than 11 seconds being the average performance for subjects over 60 years old (Bohannon, 2006; Buatois et al., 2008). The cognitive fitness of the aging subjects was established using the Mini-Mental State Examination (MMSE—(Folstein et al., 1975). All the aging subjects in the study had MSSE scores >24, indicating the absence of dementia (Lezak, 2004). All the elderly participants included in our study were considered active, practicing at least one supervised physical activity per week (e.g. gymnastics).

### 2.2. Motor task

All the participants performed a WBR movement (Figure 1-1). The experimental procedures have been used and validated in previous studies (Pozzo et al., 2002; Schmid et al., 2006; Berret et al., 2009; Chiovetto et al., 2010; Casteran et al., 2013). We asked participants to perform a WBR movement simultaneously with their two index fingers to touch two targets. The targets (4 × 2 cm) were separated by 0.5 m from their centres and positioned on a piece of wood. They were placed at distances corresponding to 30% of each participant’s height in the anteroposterior plane (AP) and in the vertical plane (V). Distances were measured from the distal end of each participant’s big toe. Participants started from an upright position. Their hands were positioned so that the hypothenar eminence was in contact with the thighs. Only, the index finger remained extended, while the rest of the fingers were bent. Target accuracy was the primary constraint imposed on the participants.

**Figure 1.**
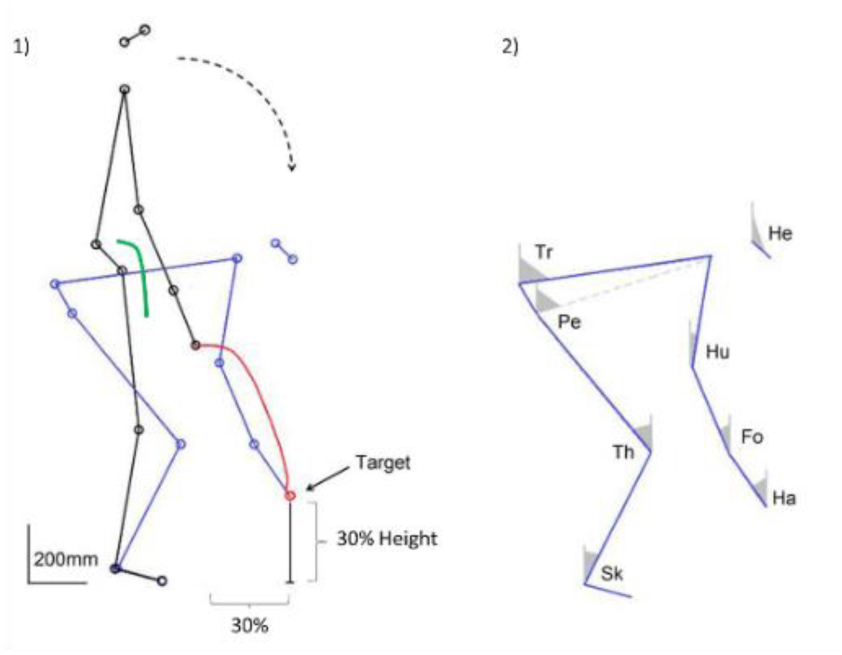
Stick diagrams. 1) Stick diagrams of a whole-body reaching movement towards a target positioned at 30% on the anteroposterior axis and on the vertical axis. The displacement of the finger marker is depicted in red, and the displacement of the CoM is depicted in green. 2) Computed elevation angles for the movements include the Shank (Sk), Thigh (Th), Pelvis (Pe), Trunk (Tr), Head (He), Humerus (Hu), Forearm (Fo) and Hand (Ha).

### 2.3. Data collection

We used an optoelectronic device (VICON, sampling frequency 200 Hz) with three cameras to capture movement kinematics in three dimensions (3D). Twelve retro-reflective markers (0.2 m in diameter) were placed at various anatomical locations on the right side of the body (external cantus of eye, auditory meatus, acromial process, humeral condyle, ulnar styloid process, apex of the index finger, L5 vertebra, greater trochanter, knee interstitial joint space, external malleolus, fifth metatarsal head of the foot, and the middle of arm) in order to have 3D with the VICON system. We used a nine-segment model similar to our previous studies of the same movement (Berret et al., 2009; Chiovetto et al., 2010; Casteran et al., 2013).

All processing of the 3D marker positions was performed with custom software written in Python in the Jupyter Notebook environment 7.0.6. Before the computation of the kinematic data, the recorded marker position signals were low-pass filtered using a fifth-order Butterworth filter at a cutoff frequency of 7.5 Hz (Berret et al., 2009) (Python 3). The filtering was followed using interpolation routines (Python 3) so that all trajectories irrespective of execution duration lay along a 200-point time base.

### 2.4. Data preparation and dataset

#### 2.4.1. Kinematic computations

Movement onset was defined as the time when the velocity of the finger exceeded 5% of its peak and movement cessation was noted likewise when this velocity dropped below the 5% threshold (Sergio and Ostry, 1994). Kinematic parameters including velocity of the finger in 3D and kinematic parameters related to angular displacements were computed using previously reported techniques (Papaxanthis et al., 2005; Berret et al., 2009; Casteran et al., 2013 and 2018) using Python 3. The following eight elevation angles (angle between the vertical and the segment) were calculated: Shank (Sk), Thigh (Th), Pelvis (Pe), Trunk (Tr), Humerus (Hu), Forearm (Fo), Hand (Ha), and Head (He) (Figure 1-2).

**Figure 2.**
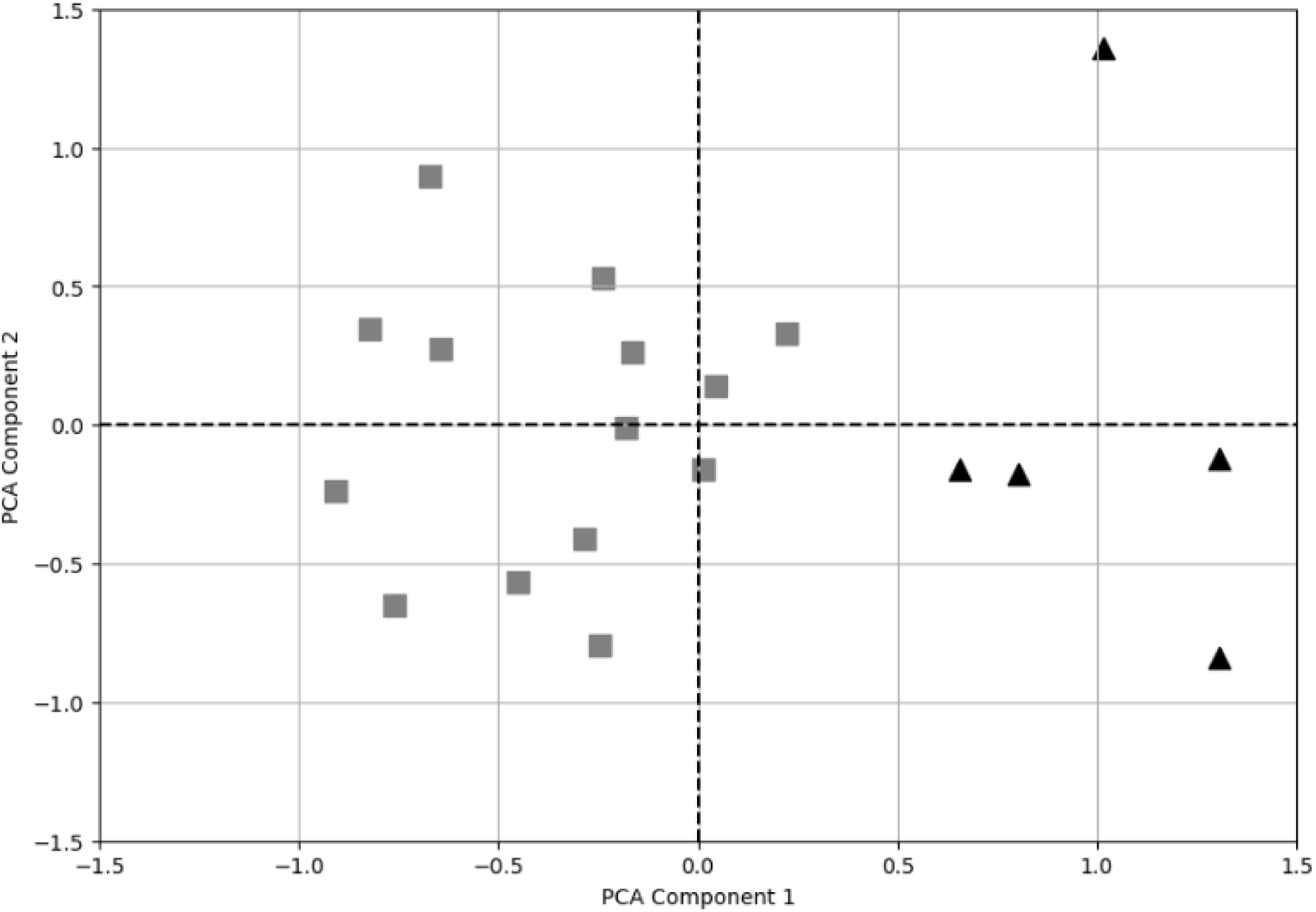
K-means clustering. This graph illustrates the two distinct motor pattern clusters, Hip-Back (HB) and Ankle-Knee (AK), based on Principal Component Analysis (PCA). The x-axis represents the first principal component, and the y-axis represents the second principal component. The Ankle-Knee (AK) cluster is depicted using light grey square markers, while the Hip-Back (HB) cluster is shown with dark grey triangle markers.

#### 2.4.2. Center of mass

We calculated the CoM displacements in 3D to characterize the way equilibrium was managed during the WBR movement (Figure 1-1 – green line). This estimation was made from an eight-segment mathematical model using rigid segments (Head, Trunk, Thigh, Shank, Foot, Upper arm, Forearm, and Hand). For this, we used the anthropometric parameters described by Winter (2009) and validated by Stapley et al. (1999) and Berret et al. (2009) in previous studies of WBR movement. Stapley et al. (1999) had compared the modeled CoM and measured center of pressure position using a force platform, during quiet stance as well as the time series of measured and estimate ground reaction forces. These studies showed that such a model provided a realistic representation of the WBR CoM position.

#### 2.4.3. Feature extraction

We extracted 21 features from the recorded data, from distinct categories. This approach allowed for a comprehensive analysis of the kinematics and dynamics involved in the movements studied, based on studies from Berret et al. (2009) and Casteran et al. (2013 and 2018).

##### Velocity kinematic parameters of finger movements

We investigated three-dimensional finger kinematics by extracting the following features: peak velocity, relative time to peak velocity and the number of velocity peaks.

##### Dynamics of center of mass displacement

The velocity profile of the CoM was analyzed in 3D and further decomposed into antero-posterior (AP) and vertical (V) components. We extracted the peak velocity, relative time to peak of velocity and number of velocity peaks from the velocity of the 3D CoM, AP CoM and V CoM. Additionally, the relative crossover point was extracted to further understand the CoM dynamics, as described in Casteran et al. (2018).

##### Angular displacement kinematics

We measured the angular excursion for eight elevation angles: head, humerus, forearm, hand, trunk, pelvis, thigh, and shank as described in Casteran et al. (2013).

#### 2.4.4. Dataset

The dataset was organized into a matrix of *n x m* where *n* represents the number of participants (19 participants) and *m* the number of features.

### 2.5. Clustering methodology

We used the K-means clustering algorithm to identify patterns in our dataset by grouping data into distinct, non-overlapping clusters (*k*) based on feature similarity. This method minimizes within-cluster variance and maximizes inter-cluster distances using Euclidean distance. Our process included data standardization, feature reduction, determining the optimal number of clusters, and executing the clustering.

The clustering was conducted using Python in the Jupyter Notebook environment 7.0.6. We used several libraries: Numpy (version 1.24.3); Pandas (version 2.0.3); Sklearn (version scikit-learn 1.3.0) including *KMeans, PCA, silhouette_score, pairwise, davies_bouldin_score, MinMaxScaler*.

#### 2.5.1. Data standardization

In our study, following the recommendation by Halilaj et al. (2018) to standardize data for improved model performance. Therefore, we used Min-Max standardization, utilizing the *MinMaxScaler* library from *Scikit-learn*, which normalizes the data to have a mean of zero and a standard deviation of one, thus facilitating comparison across different features by giving them equal weight (Pedregosa et al., 2011).

#### 2.5.2. Feature reduction: principal component analysis

A Principal Component Analysis (PCA) (Jolliffe, 2002) with two components was applied to the standardised dataset, to preserve most of the variation in the data and to reduce the dimensionality of the feature space. The standardized features (*MinMaxScaler*) and their correlation matrices were used for this computation, and the results obtained was then used for the clustering algorithm, similarly to the methodology of Beaudette et al. (2019).

#### 2.5.3. Optimal cluster determination

Identifying the ideal number of clusters, *k*, was a critical step in our methodology. We employed a data-driven strategy based on the silhouette score to find the most suitable *k*, iterating through a range of *k* values from 2 to 11 (Rousseeuw, 1987; Kaufman and Rousseeuw, 2009; Sawacha et al., 2020; Higgins et al., 2023). We used the default Lloyd algorithm, and we ran the clustering algorithm 50 times with different centroid seeds. For each *k*, we applied the K-means algorithm to the data and evaluated the silhouette score. The k value producing the best silhouette score was selected, indicating the most coherent cluster configuration. A higher silhouette score indicates better defined clusters (Rousseeuw, 1987). This process ensured the chosen number of clusters maximized intra-cluster cohesion and inter-cluster separation.

### 2.6. Clustering performances

To evaluate the K-means effectiveness, we computed several metrics providing a multifaceted view of the models’ performance:

- Silhouette score (Rousseeuw, 1987) : measures how similar each point is to its own cluster compared to other clusters. The optimal value is 1, while −1 generally indicate that a sample has been assigned to the wrong cluster. Values close to 0 suggest that the clusters overlap.
- Dunn index (Dunn, 1974): it is the ratio of the smallest distance between observations not in the same cluster to the largest intra-cluster distance. Higher values suggest well-separated clusters, indicating better clustering quality.
- Davies-Boulding score (Davies and Bouldin, 1979) : the score is a measure of cluster quality. It is defined as the average similarity measure of each cluster with its most similar cluster, where similarity is the ratio of within-cluster distances to between-cluster distances. Thus, clusters which are farther apart and less dispersed will result in a better score. The minimum score is zero, with lower values indicating better clustering.

### 2.7. Statistical analysis

To ascertain that there are no differences between young and active older adults on movement duration, independently of the cluster assignment, a two-tailed Student’s t-test was applied.

Following the identification of the most effective model, an examination of the resulting optimal K-means model will be conducted. This analysis will focus on interpreting the features of the clusters to discern motor patterns.

The cluster assignments from the K-means algorithm served as the independent variable, with the number of levels corresponding to the determined number of clusters.

Age class (older vs young adults) and gender (woman vs man) are categorical variables used as control variables to identify differences within clusters. Consequently, we applied a Fischer’s exact test.

The kinematic variables and the movement duration were considered as dependent variables in this study. Prior to conducting inferential statistical tests, assessments for normality and homoscedasticity of the data were performed to ensure the validity of the statistical assumptions. Depending on the outcomes of these preliminary tests, a two-tailed Student’s t-test was applied to examine the differences among clusters. If the assumption of equal variances was violated, we used Welch’s t-test (Delacre et al., 2017).

An alpha threshold of 0.05 has been established for statistical significance. Effect sizes for significant findings was quantified using Cohen’s d for Student’s and Welch’s t-test.

All statistical analyses were performed using the Jamovi software v2.3.28 (jamovi project (2022)).

## 3. Results

### 3.1. Performance evaluation of K-means clustering

The K-means identified *k* = 2 as the optimal number (Figure 2). The performance of the model is summarized as follows: the silhouette score, was 0.46; the Dunn index was 0.29; the Davies-Bouldin index was 0.87.

### 3.2. Analysis of optimal clustering results

#### 3.2.1. Age, gender and movement duration in clustering

Statistical analysis reveals no significant difference in gender (p = 0.33, Fisher’s exact test), in age class (young vs older adults) (p = 0.141, Fisher’s exact test) and in movement duration (p = 0.73, Student’s t test) between the two identified clusters. Furthermore, no significant difference in movement duration (p = 0.52, Student’s t test) between young vs older adults was found. Despite the absence of statistical significance in age class comparisons, descriptive analysis of the age distributions indicated distinct skewness patterns within the clusters. The HB cluster exhibited a skewness of −2.17, suggesting a tendency towards older adults. In contrast, the AK cluster demonstrated a skewness of 0.65, indicating a tendency towards younger adults. Additionally, the median ages further underscore the age distribution within each cluster, with a median age of 71 years for the HB cluster and 23.6 years for the AK cluster (see Table 1 in the supplementary material).

#### 3.2.2. Patterns of movement

The descriptive analysis of each feature in each cluster is depicted in Table 1 in the supplementary material. The homoscedasticity and normality of features were checked before applying the t-test. The two-tailed Student’s and Welch’s t-test, revealed six features exhibiting statistically significant differences between clusters: humerus angle excursion (t (17) = 2.12; p = 0.04, Cohen’s d = 1.10); trunk angle excursion (t (17) = −5.98; p <0.001, Cohen’s d = −3.11); pelvis angle excursion (t (17) = −5.04; p <0.001, Cohen’s d = −2.62); thigh angle excursion (t (17) = 7.81; p <0.001, Cohen’s d = 4.07); shank angle excursion (t (17) = 6.76; p <0.001, Cohen’s d = 3.52); maximal vertical velocity of the CoM (t (17) = 6.32; p <0.001, Cohen’s d = 2.70). All analysis are in the supplementary material (Table 2, 3, 4).

## 4. Discussion

The primary objective of this study was to determine whether K-means could identify different movement patterns based on motor control features and whether they were age-dependent or independent. Additionally, we aimed to present the contribution and advantages of unsupervised machine learning to detect motor patterns and a way of applying K-means in the field of motor control research, based on the current best practice.

### 4.1. Motor control and aging

#### Pattern identification

Our study revealed two distinct motor patterns: cluster HB exhibiting higher trunk and pelvis excursion, indicative of a hip-back strategy; cluster AK showing greater humerus, thigh and shank excursion along with increased maximal vertical velocity of the CoM, suggesting a knee-ankle strategy. The nature of these two profiles is similar of those demonstrated by numerous authors who were interested in postural displacement following reactions to an external stimulus, inducing reactive postural adjustments (Roberts and Stenhouse, 1977; Marsden et al., 1981; Horak and Nashner, 1986). These researches led to the identification of two internal strategies: “ankle strategy” and “hip strategy” (Nashner, 1985; Nashner and McCollum, 1985). In this type of paradigm, it has been shown that the response to postural perturbations changes with age (Afschrift et al., 2018). The shift from an ankle-centred to a hip-centred strategy with increasing magnitude of perturbation occurs at lower accelerations in older adults than in younger adults (Horak and Nashner, 1986; Fujimoto et al., 2013; Hsu et al., 2013). Conventional analysis in motor control research focuses mainly on differences between groups (i.e. young vs older adults). In comparison, our study using unsupervised machine learning based on motor control features and not an a priori grouping, showed age independent motor patterns. This approach provides a complementary analysis as highlighted by Ackermans et al. (2019). Their work, using K-means clustering, demonstrated the limitations of conventional comparisons of biomechanical parameters in capturing the complexity of fall risk. They encourage a multivariate approach to go beyond conventional group comparisons. Overall, this approach opens the possibility of profiling individuals according to their motor patterns instead of demographic characteristics usually used to define groups.

#### Age-related and non-age-related patterns

While not focusing on whole-body reaching movement, previous studies have demonstrated that older adults : (i) exhibited a reduced stability limit for their CoM control in the forward–backward direction during upright standing (Fujimoto et al., 2013; Hernandez et al., 2013); (ii) decreased the movement of the center of pressure toward the backward direction during gait initiation from an upright standing position than younger individuals (Halliday et al., 1998; Khanmohammadi et al., 2015); (iii) may moving their CoM forward by increasing the trunk flexion angle during deep-squat movement to avoid falling backward (Honda et al., 2022). In the study of a whole-body reaching movement, Casteran et al. (2018), through CoM analysis, suggest an age-related adaptation prioritising balance control over the minimization of absolute work. Therefore, we could expect that the cluster HB, found in our study could be age-dependent and corresponds to older adults and the cluster AK to young adults. Nevertheless, in our study, motor patterns are age independent. In fact, based on the age class (young vs elderly), there is not statistical difference between the two clusters. However, it should be noted that the median values of the age and the value of skewness of the age class suggest a tendency for HB to be associated with older adults, whereas AK tends to correspond to younger adults. The trend towards a hip-back strategy in older adults is also echoed in the work of Papa & Cappozzo (2000), who noted increased trunk flexion during the sit-to-stand task in older adults.

### 4.2. Clustering methodology and assessment

#### Number of clusters

The K-means algorithm requires a predefined number of clusters (*k*), which is often unknown and can significantly affects the algorithm’s performance. Identifying the optimal *k* is crucial, and Pham et al. (2005) describe methods like user choices, statistical criteria, and visual aids, each with specific trade-offs depending on data characteristics and model assumptions. In our study, influenced by findings from Mulyani et al. (2023), Sai et al. (2017), Shahapure & Nicholas (2020), which underscored the silhouette score’s ability to enhance clustering quality, we propose to use it to determine the optimal *k*.

#### Feature reduction

Given the curse of dimensionality, it is advisable to use feature reduction to identify relevant features and thus reduce dimensions (Jovic et al., 2015). In K-means studies specifically, Zakharov (2016) shows that the most common way to dimension reduction is by applying PCA or the Exploratory Factor Analysis (EFA). Halilaj et al. (2018) emphasized also the utility of automatic feature extraction methods like PCA for dimensionality reduction while retaining important variations. In our case, we could therefore reduce the dataset from 21 features to 2 principal component and capture the more relevant data, thereby improving clustering performances.

#### Model performance

Our study’s findings demonstrate clustering performance through several metrics. The silhouette score indicates well-separated clusters with closely grouped points within each cluster. The Dunn Index confirms reasonable cluster separation, while the Davies-Bouldin Index shows that the clusters are compact with low intra-cluster dispersion and significant inter-cluster distance. Combined, these measures suggest that our clustering model is strong and effective when compared to previous motor control research (Higgins et al. (2022)).

### 4.3. Limitations and future perspectives

This study provides insights into clustering methodologies for motor control analyses, based on kinematic data. However, several limitations can be highlighted. First, the population sample was limited to young and older active adults. Second, the study exclusively used kinematic data for movement profile identification. Integrating other data modalities, such as kinetic and electromyographical data, and clinical assessments like strength and functional tests, could provide a more comprehensive approach to movement dynamics and contribute to a more detailed and accurate profiling. Given these limitations, we encourage future studies to investigate these areas. We also suggest employing unsupervised machine learning to complement conventional analysis, in order to investigate the complexity of motor control. While our study focused on natural speed, analysing different speed conditions would enable to study motor control adaptation across all ages, at faster or slower speeds.

## 5. Conclusion

In conclusion, this study proposes to identify motor patterns based on motor control features using the K-means algorithm. This method integrates the latest recommendation, including data standardization, feature reduction and determination of the *k* value of clustering using the silhouette score. We could identify two distinct age-independent movement patterns based on kinematic data, moving beyond conventional age-based grouping. The first profile has higher humerus, ankle and knee angular excursions, along with higher vertical velocity of CoM. The second has higher hip and back angular excursions, along with lower vertical velocity CoM. The clustering methodology demonstrated its effectiveness to identify distinct motor patterns based solely on motor control features independently of age-grouping. We propose to include analyses based on motor profiles complementary to demographic characteristics of participants. This combination could facilitate the identification of motor disorder profiles. For example, it could help identify profiles related to joint and muscle conditions, fall risk, loss of autonomy, deconditioning, and osteoarthritis.

## Acknowledgments

Michel Pfaff was supported by a doctoral grant from the Université de Lorraine. The funders had no role in the study design, data collection and analysis, decision to publish or preparation of the manuscript.

