## Supplementary material for "Identification of whole-body reaching movement phenotypes in young and older active adults: an unsupervised machine learning approach"

**Table 1.**

**Descriptive statistics of each cluster**

| | | | | | | | | **Skewness** | |
|---|---|---|---|---|---|---|---|---|---|
| | **Cluster** | **N** | **Mean** | **Median** | **SD** | **Minimum** | **Maximum** | **Skewness** | **SES** |
| He_Excursion | AK | 14 | 28.31 | 30.929 | 9.935 | 15.147 | 48.289 | 0.2405 | 0.6 |
| | HB | 5 | 27.73 | 30.671 | 12.304 | 12.933 | 39.9 | -0.3401 | 0.91 |
| Hu_Excursion | AK | | 15.94 | 15.683 | 3.9128 | 8.77 | 21.145 | -0.2309 | 0.6 |
| | HB | | 11.46 | 12.712 | 4.4212 | 5.462 | 16.961 | -0.2834 | 0.91 |
| Fo_Excursion | AK | | 32.19 | 31.895 | 7.7799 | 19.565 | 49.442 | 0.4699 | 0.6 |
| | HB | | 29.95 | 35.109 | 9.5983 | 17.612 | 38.016 | -0.651 | 0.91 |
| Ha_Excursion | AK | | 42.03 | 41.716 | 9.9095 | 29.027 | 60.623 | 0.4222 | 0.6 |
| | HB | | 31.93 | 29.934 | 15.208 | 10.21 | 52.196 | -0.2069 | 0.91 |
| Tr_Excursion | AK | | 40.2 | 39.539 | 5.1246 | 33.788 | 48.482 | 0.3826 | 0.6 |
| | HB | | 55.93 | 56.856 | 4.7787 | 50.626 | 62.857 | 0.5208 | 0.91 |
| Pe_Excursion | AK | | 43.82 | 42.198 | 7.6236 | 32.167 | 57.022 | 0.3518 | 0.6 |
| | HB | | 63.05 | 63.697 | 6.1809 | 54.184 | 68.791 | -0.64 | 0.91 |
| Th_Excursion | AK | | 39.96 | 41.186 | 6.4761 | 28.322 | 48.904 | -0.4229 | 0.6 |
| | HB | | 14.1 | 13.001 | 5.9199 | 5.786 | 21.528 | -0.254 | 0.91 |
| Sk_Excursion | AK | | 22.45 | 23.257 | 6.0456 | 11.191 | 31.074 | -0.6232 | 0.6 |
| | HB | | 3.644 | 3.294 | 1.5548 | 2.003 | 5.812 | 0.572 | 0.91 |
| Cross | AK | | 22.12 | 24.167 | 8.2212 | 6.313 | 37.576 | -0.3669 | 0.6 |
| | HB | | 25.93 | 27.626 | 8.1277 | 12.346 | 33.283 | -1.55 | 0.91 |
| V_CoM_V_Max | AK | | 2.696 | 2.705 | 0.3462 | 2.202 | 3.152 | -0.1585 | 0.6 |
| | HB | | 1.977 | 1.958 | 0.1477 | 1.827 | 2.212 | 1.1176 | 0.91 |
| V_CoM_V_RTP | AK | | 0.548 | 0.544 | 0.0663 | 0.417 | 0.633 | -0.5884 | 0.6 |
| | HB | | 0.54 | 0.533 | 0.0593 | 0.471 | 0.613 | 0.1438 | 0.91 |
| V_CoM_V_Nb_Peaks | AK | | 2.872 | 2.75 | 1.0173 | 1.333 | 5.125 | 0.5578 | 0.6 |
| | HB | | 3.029 | 2.7 | 1.477 | 1.444 | 5.1 | 0.5745 | 0.91 |
| V_CoM_AP_Max | AK | | 0.752 | 0.665 | 0.2645 | 0.433 | 1.457 | 1.552 | 0.6 |
| | HB | | 0.77 | 0.77 | 0.0818 | 0.641 | 0.848 | -1.0928 | 0.91 |
| V_CoM_AP_RTP | AK | | 0.371 | 0.363 | 0.102 | 0.202 | 0.593 | 0.5021 | 0.6 |
| | HB | | 0.397 | 0.395 | 0.1338 | 0.186 | 0.521 | -1.0781 | 0.91 |
| V_CoM_AP_Nb_Peaks | AK | | 6.352 | 5.85 | 1.8763 | 3.3 | 9.667 | 0.4593 | 0.6 |
| | HB | | 5.391 | 5.4 | 1.7588 | 2.778 | 7.2 | -0.7252 | 0.91 |
| V_F_3D_Max | AK | | 0.634 | 0.65 | 0.1679 | 0.394 | 0.933 | 0.1034 | 0.6 |
| | HB | | 0.754 | 0.713 | 0.4085 | 0.161 | 1.139 | -0.5976 | 0.91 |
| V_F_3D_RTP | AK | | 0.371 | 0.397 | 0.082 | 0.225 | 0.491 | -0.5455 | 0.6 |
| | HB | | 0.358 | 0.358 | 0.0756 | 0.247 | 0.458 | -0.3891 | 0.91 |
| V_F_3D_Nb_Peaks | AK | | 2.961 | 2.55 | 0.9722 | 2 | 5.222 | 1.3285 | 0.6 |
| | HB | | 3.056 | 2.222 | 1.6903 | 1.7 | 5.8 | 1.428 | 0.91 |
| V_CoM_3D_Max | AK | | 0.404 | 0.404 | 0.0791 | 0.286 | 0.534 | 0.1637 | 0.6 |
| | HB | | 0.324 | 0.322 | 0.0802 | 0.203 | 0.416 | -0.7081 | 0.91 |
| V_CoM_3D_RTP | AK | | 0.492 | 0.488 | 0.0627 | 0.389 | 0.613 | 0.0422 | 0.6 |
| | HB | | 0.468 | 0.461 | 0.0319 | 0.434 | 0.509 | 0.3701 | 0.91 |

| | | | | | | | | | |
|---|---|---|---|---|---|---|---|---|---|
| V_CoM_3D_Nb_Peaks | AK | | 3.236 | 3.2 | 1.1873 | 1.8 | 5.3 | 0.5583 | 0.6 |
| | HB | | 4.384 | 3.778 | 2.0482 | 2.444 | 7.8 | 1.5212 | 0.91 |
| Age | AK | | 40.64 | 25.5 | 23.562 | 20 | 72 | 0.6515 | 0.6 |
| | HB | | 62 | 71 | 20.236 | 26 | 74 | -2.1749 | 0.91 |
| MD | AK | | 1.414 | 1.325 | 0.2576 | 1.079 | 1.828 | 0.3848 | 0.6 |
| | HB | | 1.367 | 1.393 | 0.4234 | 0.962 | 2.037 | 1.101 | 0.91 |

Descriptive analysis of kinematic features for clusters (Ankle-Knee (AK) and Hip-Back (HB)), alongside control variable: age and movement duration (MD). The features analysed include the angle excursions (_Excursion) of different parts of the body (Head (He), Humerus (Hu), Forearm (Fo), Hand (Ha), Trunk (Tr), Pelvis (Pe), Thigh (Th), Shank (Sk), the cross-over point (Cross) and several parameters measuring the velocity of the centre of mass (V_CoM) differentiated into vertical (_V), anterior-posterior (_AP) and 3D movement for the CoM and the velocity of the fingers in 3D (V_F_3D). These velocity measurements include maximum velocity (_Max), relative time to reach maximum velocity (_RTP) and number of velocity peaks (_Nb_Peaks). This table includes the following statistical metrics for each feature: sample size (N), mean, median, standard deviation (SD), minimum values, maximum values, Skewness score and Standard Error of Skewness (SES).

**Table 2.**

**Homogeneity of variances tests (Levene's)**

| | F | df | df2 | p |
|---|---|---|---|---|
| Hu_Excursion | 0.0629 | 1 | 17 | 0.805 |
| Tr_Excursion | 0.4271 | 1 | 17 | 0.522 |
| Pe_Excursion | 0.4001 | 1 | 17 | 0.535 |
| Th_Excursion | 0.2874 | 1 | 17 | 0.599 |
| Sk_Excursion | 3.8113 | 1 | 17 | 0.068 |
| V_CoM_V_Max | 6.2342 | 1 | 17 | 0.023 |

Results from the Levene's test for six features identified with significant differences between clusters (AK vs HB): the angle excursion (_Excursion) of the Humerus (Hu), Trunk (Tr), Pelvis (Pe), Thigh (Th), Shank (Sk), and the maximal vertical velocity of the centre of mass (V_CoM_V_Max). Statistical metrics including F values, degrees of freedom 1 (df1), degrees of freedom 2 (df2) and p value for each feature comparison are presented.

**Table 3.**

**Test of normality (Shapiro-Wilk)**

| | W | p |
|---|---|---|
| Hu_Excursion | 0.955 | 0.482 |
| Tr_Excursion | 0.916 | 0.097 |
| Pe_Excursion | 0.965 | 0.683 |
| Th_Excursion | 0.953 | 0.450 |
| Sk_Excursion | 0.929 | 0.168 |
| V_CoM_V_Max | 0.948 | 0.372 |

Results from the test of normality (Shapiro-Wilk) for six features identified with significant differences between clusters (AK vs HB): the angle excursion (_Excursion) of the Humerus (Hu), Trunk (Tr), Pelvis (Pe), Thigh (Th), Shank (Sk), and the maximal vertical velocity of the centre of mass (V_CoM_V_Max). Statistical metrics including W values and p-values for each feature comparison are presented.

**Table 4.**

**Independent samples t-test comparisons between clusters**

| | | Statistic | df | p | | Effect Size |
|---|---|---|---|---|---|---|
| Hu_Excursion | Student's t | 2.13 | 17.0 | 0.048 | Cohen's d | 1.11 |
| Tr_Excursion | Student's t | -5.99 | 17.0 | < .001 | Cohen's d | -3.12 |
| Pe_Excursion | Student's t | -5.05 | 17.0 | < .001 | Cohen's d | -2.63 |
| Th_Excursion | Student's t | 7.82 | 17.0 | < .001 | Cohen's d | 4.07 |
| Sk_Excursion | Student's t | 6.76 | 17.0 | < .001 | Cohen's d | 3.52 |
| V_CoM_V_Max | Welch's t | 6.32 | 16.06 | < .001 | Cohen's d | 2.70 |

Results from the independent samples t-test and Welch's test for six features identified with significant differences between clusters ($H_a$ μ HB ≠ μ AK): the angle excursion (_Excursion) of the Humerus (Hu), Trunk (Tr), Pelvis (Pe), Thigh (Th), Shank (Sk), and the maximal vertical velocity of the centre of mass (V_CoM_V_Max). Statistical metrics including Student's and Welch's t-test values, degrees of freedom (df), p-values, mean difference, and Cohen's d effect sizes for each feature comparison are presented.
